# A bidirectional *nanAKE* locus enables sialic acid catabolism in gut microbiome member *Hungatella hathewayi*

**DOI:** 10.64898/2026.05.18.725967

**Authors:** Vienvilay Phandanouvong-Lozano, Letizia Pastore, Gabriel Miller, Kathryn Y. Lin, Ashley R. Wolf

**Affiliations:** School of Public Health Division of Infectious Diseases and Vaccinology - University of California Berkeley, Berkeley, CA; Department of Integrative Biology - University of California Berkeley, Berkeley, CA; Department of Molecular and Cell Biology - University of California Berkeley, Berkeley, CA; Center of Computational Biology - University of California Berkeley, Berkeley, CA

## Abstract

Sialic acids are abundant components of host- and diet-derived glycans in the human gut and serve as important nutrients that shape microbial fitness and interspecies competition. Excess free sialic acids are also linked to inflammation and pathogen susceptibility. While well-studied gut bacteria such as *E. coli* and *Bacteroides* spp. catabolize sialic acids via the NanAKE or NanLE-RokA pathways, the metabolic capacity of many microbiome members remains undefined. To identify sialic acid catabolizing bacteria, we cultured fecal samples from healthy human donors. The gut anaerobe *Hungatella hathewayi* was selected under sialic acid-supplemented conditions. *H. hathewayi* is a poorly characterized gram-positive *Lachnospiraceae* associated with long-lived individuals and purine metabolism. Here we establish that *H. hathewayi grows* robustly on sialic acids as a sole carbon source using a pathway homologous to the canonical NanAKE system of *E. coli*, despite the species’ phylogenetic distance. We functionally validated these orthologs through growth assays and heterologous complementation in *E. coli* knockout strains. Comparative analyses further showed that key catalytic residues in *H. hathewayi* NanA are conserved despite overall sequence divergence from *E. coli*. Additionally, we find that colocalized sialic acid transporters and regulatory proteins are not orthologous to *E. coli* proteins and instead are related to proteins from other gut anaerobes. Together, these findings expand our understanding of sialic acid utilization within the human gut microbiome. We identify *H. hathewayi* as an overlooked but capable sialic acid degrader that can contribute to modulation of gut sialic acid levels and related inflammation.

**Importance:** Sialic acids play an important role in mammalian and microbial signalling. Excess free sialic acids increase susceptibility to gut pathogens and induce inflammation. Gut bacteria can both generate and consume free sialic acids, and these pathways are conserved across diverse bacteria. *E. coli* and *B. fragilis* consume sialic acids as a carbon source, decreasing free sialic acid levels. We identify *H. hathewayi* as another bacteria capable of sialic acid consumption and define the enzymes responsible. *H. hathewayi* is a prevalent member of the human gut microbiome, but it is not genetically tractable, limiting enzymatic characterization. *H. hathewayi* is enriched in the gut microbiomes of long-lived individuals and expected to be an important contributor to purine degradation to limit gout risk. Defining sialic acid catabolism in non-model species is essential to understanding the evolution and conservation of this pathway as well as how nutrient competition shapes gut microbiome composition.

## Introduction

The sialic acid family of monosaccharides is a diverse set of nine-carbon sugars synthesized across the tree of life. Sialic acids often serve as terminal residues of cell surface glycoconjugates in gut bacteria and mammals, and they are abundant in the mammalian gut where they can fuel microbiota metabolism (1–3). Sialic acid variants exhibit remarkable structural diversity, with more than fifty naturally occurring chemical modifications such hydroxylation, *O*-acetylation, and *O*-sulfation (1, 2, 4, 5). This diversity underlies their involvement in a wide range of physiological processes including cellular metabolism, immune modulation, development, and host–microbiota interactions (2, 6, 7). In humans, *N*-acetylneuraminic acid (Neu5Ac) is the primary sialic acid variant. Neu5Ac populates the terminal ends of mucin glycan chains that constitute the carbohydrate-rich mucosal surfaces of the gastrointestinal tract, oral cavity, and reproductive tract (8–11). They are also present in specific human milk oligosaccharides (HMOs), where they contribute to neonatal gut and immune development (12–14). *N*-glycolylneuraminic acid (Neu5Gc) is not present in human tissues but is found in non-human mammals and accessible to gut microbes after meat consumption (15).

Certain gut commensal and pathogenic bacteria can utilize sialic acids found in intestinal mucus as carbon and nitrogen sources (16–19). For pathogenic bacteria, the ability to consume, acquire, or mimic host-derived sialic acids is a key factor in colonization and immune evasion (20–22). While some gut bacteria encode sialidases that directly cleave sialic acids from host mucins, many others lack sialidase activity and instead rely on free sialic acids generated by neighboring microbes (17, 23, 24). The liberation of sialic acids from mucins by sialidases in the gut environment promotes the expansion of pathogens, including *Salmonella enterica* serovar *Typhimurium* and *Clostridioides difficile*, which exploit these freed sialic acids via dedicated sialic acid catabolic pathways (25). Excess free sialic acids can also boost levels of *E. coli* and contribute to intestinal inflammation (26).

Catabolism of host- and diet-derived sialic acids provides gut bacteria with a metabolic advantage, yet the enzymes and pathways underlying this process have been characterized in few model species (22, 24, 27–29) (Fig. 1). In *Escherichia coli*, sialic acid utilization is mediated by the Nan system, in which the transcriptional regulator NanR senses Neu5Ac and induces expression of genes encoding transport and catabolism (30, 31). Free sialic acid is imported via the outer membrane porin NanC and inner membrane transporter NanT, then cleaved by *N-*acetylneuraminate lyase (NanA) into N-acetylmannosamine (ManNAc) and pyruvate. ManNAc is phosphorylated by *N-*acetylmannosamine kinase (NanK) and epimerized by *N-*acetylmannosamine-6-phosphate epimerase (NanE) to generate N-acetylglucosamine-6-phosphate (GlcNAc-6-P), which feeds into central amino sugar and glycolytic metabolism (29, 32–34). This canonical system is not restricted to Gram-negative models such as *E. coli;* Gram-positive *Bacillota* (formerly *Firmicutes*) such as *Clostridium perfringes* and *Clostridiodes difficile* have shown that sialic acid catabolism can also be mediated by a Nan system, demonstrating conservation across phylogenetically diverse bacteria (20, 28, 35).

**Figure 1.**
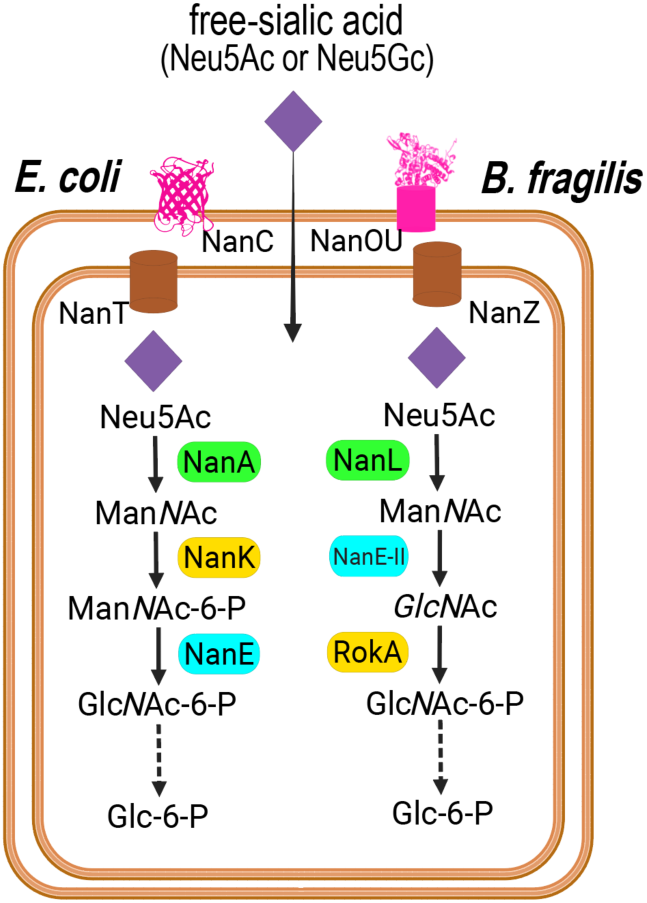
Schematic of sialic acid catabolism in *Escherichia coli* and *Bacteroides fragilis* via NanAKE and NanLEK pathway, respectively. Free-sialic acid as purple diamond, outer-membrane transporters in pink (NanC, NanOU), inner-membrane transporters in brown (NanT, NanZ), lyases in green, kinases in yellow, and epimerases in blue (NanA, NanL: *N*-acetylneuraminate lyase, NanK: *N*-acetylmannosamine kinase, NanE: *N*-acetylmannosamine-6-phosphate 2-epimerase, NanE-II: *N*-acylglucosamine 2-epimerase, RokA: ROK family sugar kinase).

While this Nan pathway represents the classical model, gut bacteria exhibit substantial diversity in how they acquire sialic acids. Some employ TonB-dependent systems, such as NanOU in Bacteroides, whereas others use ABC-type importers common among *Bacillota* (36–38). Variations in the order of enzymatic steps also occur, as seen in *Bacteroides fragilis* where an epimerase acts before a kinase (27, 39) (Fig. 1), but these pathways converge on the same GlcNAc-6-P intermediate that enters central carbon metabolism. Collectively, these systems enable bacteria to scavenge sialic acids released from host mucins and glycoproteins, linking host glycan turnover to microbial energy metabolism and colonization (18, 21, 40).

*Hungatella hathewayi* is a largely uncharacterized gut bacterium, whose genomic potential, ecological distribution, and clinical associations point to a role in host glycan utilization. *H. hathewayi* is an obligate anaerobe belonging to the phylum *Bacillota*, and was initially isolated from a chemostat inoculated with human faeces (41, 42). There are few functional studies of *H. hathewayi* isolates and it is under-represented in functional annotations (43). Genomic analyses have shown that *H. hathewayi* harbors genes consistent with saccharolytic metabolism and glycan utilization, suggesting adaptation to mucosal or host-glycan niches (43, 44). *H. hathewayi* is enriched in the gut microbiomes of long-lived individuals (45). In contrast, *H. hathewayi* has also been linked to epithelial promoter hypermethylation, a process linked to colorectal cancer (46). Although generally considered a gut commensal, *H. hathewayi* has also been implicated in rare but serious bacteremia and opportunistic infections (47–49). Taken together, these observations suggest that *H. hathewayi* occupies a distinctive niche within the gut microbiome potentially adapted for glycan-derived substrate use, but its specific metabolic role and inter-microbial interactions remain to be defined. We investigated the genetic basis and metabolic capacity for sialic acid catabolism in *H. hathewayi* to better understand how this gut commensal exploits host-derived glycans and what this may reveal about its ecological role in the intestinal environment. Microbes that can consume sialic acids without prompting inflammation may be useful as probiotics to control expansion of sialic acid consuming pathogens and pathobionts.

## Results

### Enrichment approach identifies Hungatella as a putative sialic acid consumer

To identify bacteria capable of utilizing sialic acids, we used an enrichment culturing approach with fecal suspensions from seven healthy human donors (Fig. 2A). Each sample was inoculated into base anaerobic medium (BAM, Table S1) supplemented with either *N*-acetylneuraminic acid (Neu5Ac), *N*-glycolylneuraminic acid (Neu5Gc), or porcine gastric mucin as the sole carbon source. Following anaerobic incubation, bacterial enrichment was determined via 16S rRNA gene amplicon sequencing. We identified bacterial taxa that were significantly enriched compared to baseline on specific carbon sources using ANCOM-BC (Fig. 2B). These enriched taxa are likely to be utilizers of the sialic acid or mucin substrate included in the culture.

**Figure 2.**
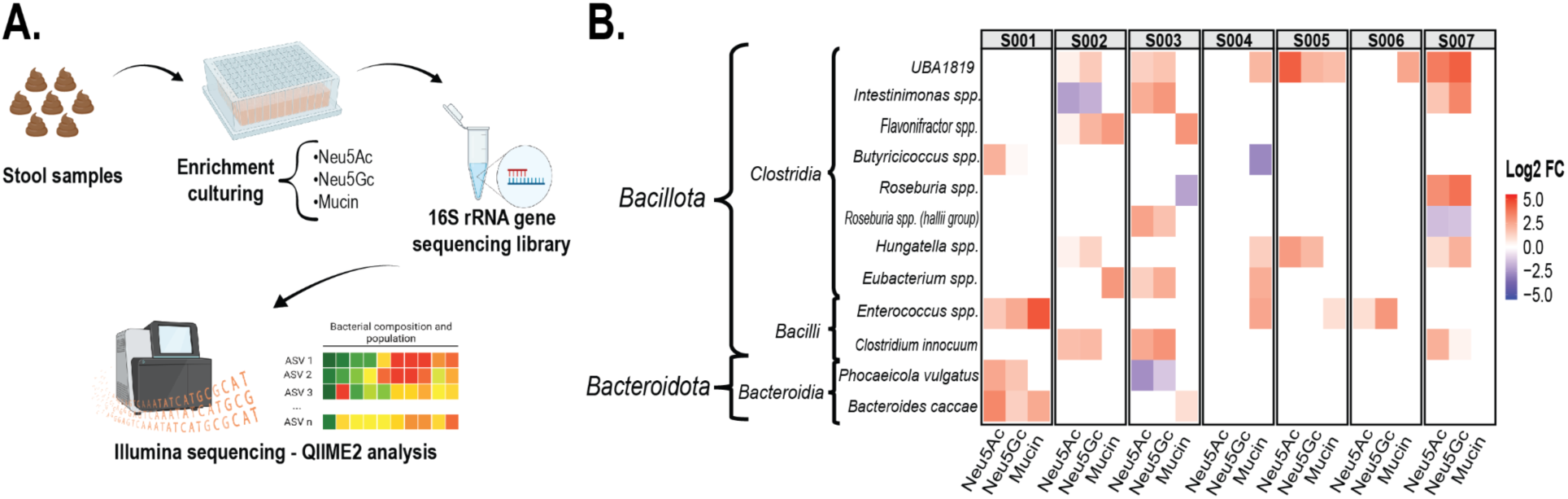
**A.** Human fecal suspensions were enriched with a defined base anaerobic medium (BAM) containing sialic acid (10 mM of Neu5Ac or Neu5Gc) or mucin (1%). Bacterial composition and abundance were determined via 16S rRNA gene sequencing. **B.** Heatmap of bacterial taxa exhibiting significant differential abundance in response to growth on the specified substrate (NeuAGc, Neu5Gc, or Mucin). Differential abundance was calculated by comparing bacterial communities from enrichment cultures against their corresponding unenriched fecal inocula (i.e., baseline microbiota prior to incubation). ANCOM-BC analysis using thresholds of Fold Change (FC) ≥2 and adjusted p-value <0.05. Samples S001–S007 represent individual donors.

We identified 13 bacterial taxa across the seven donors that were enriched under at least one condition. These taxa were primarily from the phyla *Bacillota* and *Bacteroidota*, including known sialic acid degrader *Phocaeicola vulgatus*. While enrichment patterns varied between samples, a bacterial taxon belonging to the genus *Hungatella* was enriched on growth of both sialic acids in samples from three donors (S002, S005, and S007 in Fig. 2B). This led us to hypothesize that *Hungatella* species are capable of consuming both Neu5Ac and Neu5Gc.

### *Hungatella hathewayi* genomes encode a conserved *nanAKE* cluster for sialic acid catabolism

To identify putative sialic acid-degrading genes in *Hungatella hathewayi,* we selected a total of 17 genomes, comprising the type strain DSM 13479 and 16 publicly available genomes from gut isolates with adequate genome-quality (completeness ≥ 80%, contamination ≤ 10%, and fine consistency ≥ 87%, Bacterial and Viral Bioinformatics Resource Center, Table S2). Each genome was queried for sialic acid catabolic genes using BLASTP searches against the characterized *E. coli* NanA, NanK, and NanE protein sequences, which encode enzymes capable of sialic acid degradation (29, 33). The majority of the *H. hathewayi* genomes (12 of 16), as well as the type strain DSM13479, encode a complete *nanAKE*-like cluster (Fig. S1).

We next performed comparative genomic analysis with a representative *H. hathewayi nanAKE*-like cluster and known sialic acid gene utilization clusters from *E. coli*, *Clostridioides difficile*, *Bacteroides fragilis* and *Phocaeicola vulgatus* (Fig. 3). The *H. hathewayi* locus differs from these canonical models in a few ways. In *E. coli*, the sialic acid catabolic genes form the co-directional operon *nanATEK*, accompanied by the sialic acid inner membrane transporter *nanT,* which is proton symporter from the Major Facilitator Superfamily (MFS); and regulated by *nanR*, a GntR-family transcriptional repressor positioned divergently upstream (Figs. 1 and 3) (29, 30, 34, 50, 51). In contrast, the *H. hathewayi* locus is bidirectional, with *nanK* and *nanE* being transcribed oppositely from *nanA*. Unlike the characterized systems shown in Fig. 3, *H. hathewayi* does not contain a colocalized *nanT* homolog and instead contains a nearby predicted ABC transporter system. Instead of a nanR-gntR homolog, both *H. hathewayi* and *C. difficile* encode RpiR-family transcriptional regulators (Fig. 3, Fig. S1). RpiR-family regulators have been linked to sialic acid metabolism in diverse bacteria, including *Clostridium perfringes* (31), *Haemophilus influenzae* (52, 53), *Vibrio vulnificus* (54), and *Staphylococcus aureus* (55).

**Figure 3.**
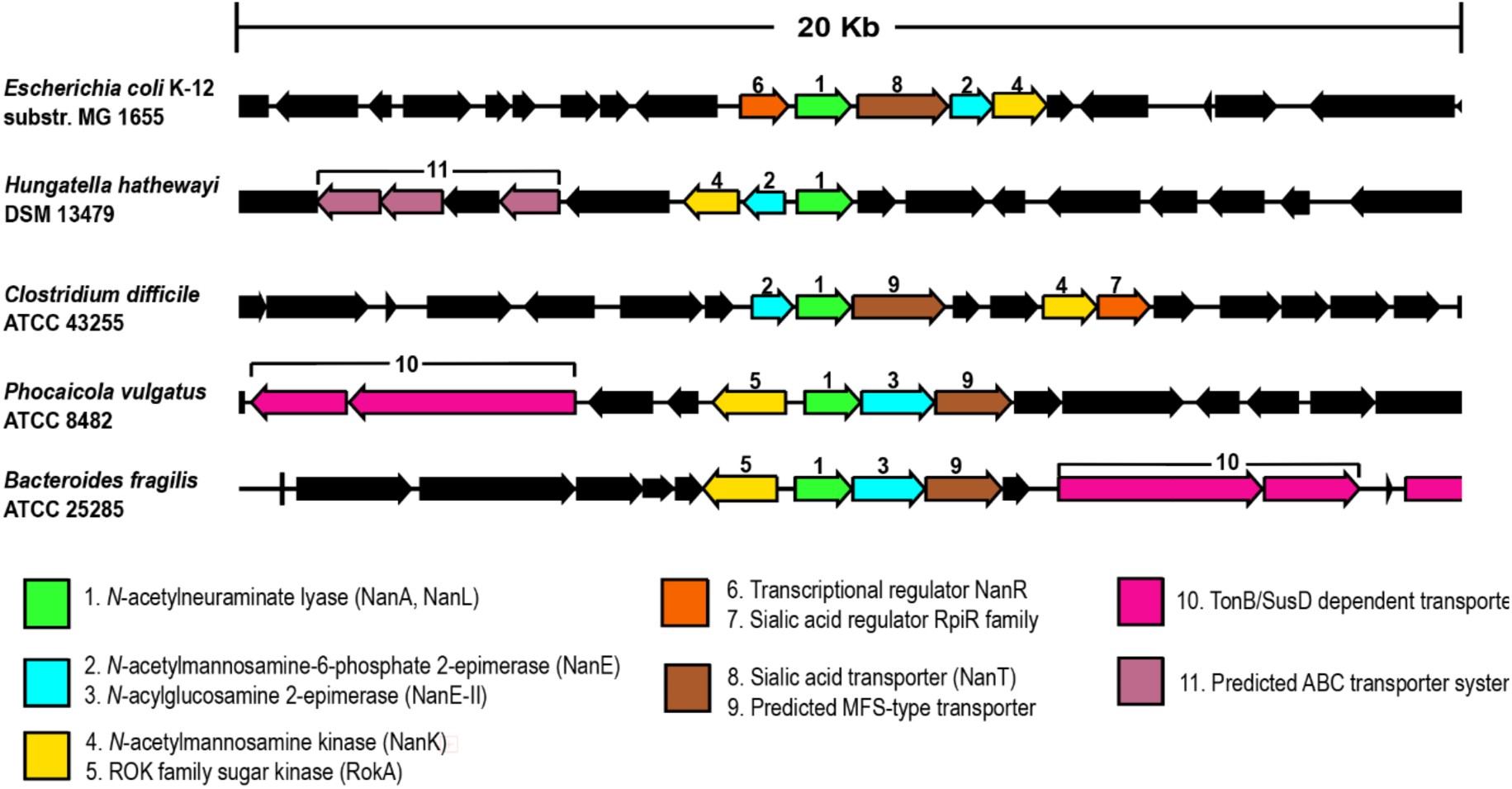
Sialic acid-catabolic genes identified in the reference strains *E. coli* K-12, *C. difficile* ATCC 43255, *B. fragilis* ATCC 25285, and *P. vulgatus* ATCC 8482 including the putative annotated genes in *H. hathewayi* DSM 13479. Genes are represented as arrows indicating orientation and scaled to their relative lengths.

*Bacteroidota* sialic acid catabolism genes are homologous yet distinct from the *E. coli* genes, as their encoded enzymes enable a slightly different breakdown pathway characterized in *B. fragilis* (27, 39) (Fig. 1). *Bacteroidota* species (e.g. *B. fragilis* and *P. vulgatus*), harbor *nan*-like genes embedded in polysaccharide utilization loci (PULs), with adjacent TonB-dependent SusC/SusD-like transporters and regulatory genes (36, 39). Based on homology, we expected that the *H. hathewayi* enzymatic breakdown of sialic acids was more likely to mimic *E. coli* than *B. fragilis*, and next sought to confirm this experimentally.

### Sialic acids support robust growth in Hungatella hathewayi

We next assessed whether *H. hathewayi* DSM13479 could utilize sialic acids as sole carbon source to confirm the functionality of its predicted catabolic pathway. Growth assays were conducted in defined base anaerobic medium (BAM) supplemented with either Neu5Ac or Neu5Gc, with glucose as a positive control, and unsupplemented BAM as a negative control. *H. hathewayi* grew robustly on both Neu5Ac and Neu5Gc, reaching higher saturation densities and higher log-phase growth rates than in glucose-containing media, while showing similar growth behavior between the two types of sialic acids (Fig. 4A, Table S3). In contrast, *E. coli* reached higher density and growth rate in glucose than in either sialic acid condition, and displayed a clear preference for Neu5Ac over Neu5Gc, with Neu5Gc supporting slower growth and prolonged lag phase. (Fig. 4B, Table S3). These results indicate that although both bacteria were capable of utilizing Neu5Ac and Neu5Gc as sole carbon sources, they exhibited distinct growth dynamics. While *E. coli* exhibits substrate-dependent differences in both growth rate and lag phase, *H. hathewayi* displayed comparable growth kinetics on both sialic acids, despite an extended lag period across all substrates.

**Figure 4.**
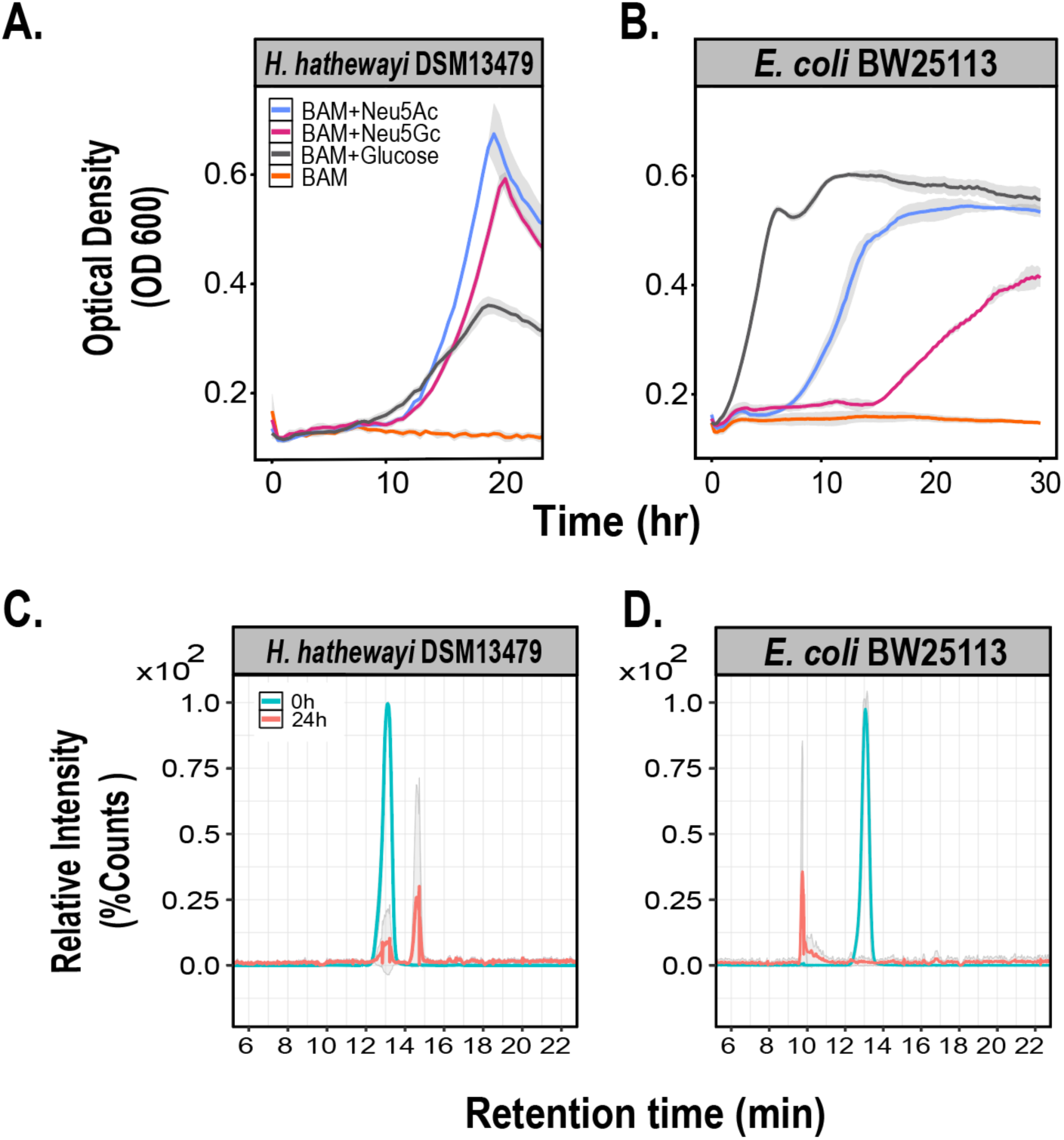
**A.** Anaerobic growth of *Hungatella hatheway*i DSM13479 and **B.** *Escherichia coli* BW25113 in defined base anaerobic media (BAM) plus the indicated sialic acid (10mM Neu5Ac or Neu5Gc). [mean of 3 replicates +/- standard deviation]. Chromatograms of culture supernatants at 0 h (blue) and 24 h (red) showing Neu5Ac depletion by **C.** *H. hatheway*i DSM13479 and **D.** *E. coli* BW25113.

Further evidence for sialic acid utilization was obtained by quantifying Neu5Ac in bacterial culture supernatants. Neu5Ac detection was based on multiple reaction monitoring (MRM) with a characteristic retention time of approximately 13.2 minutes (Fig. S2A). As described in previous studies (56, 57), fragmentation of the precursor ion (m/z 308.1) produced the expected product ions m/z 87.0, m/z 98.1, and m/z 170.0, with m/z 87.0 showing the highest intensity and therefore was selected for quantification (Fig. S2B). Quantification of Neu5Ac from culture supernatants from *H. hathewayi* and *E. coli* demonstrated complete depletion of the sialic acid from the medium by 24 hours, when both species were at stationary phase (Figs. 4C and 4D). These findings provide direct biochemical evidence that *H. hathewayi* efficiently consumes Neu5Ac during growth, consistent with the functional prediction based on the putative *nanAKE*-like catabolic pathway.

### Cross-species complementation confirms H. hathewayi sialic acid gene function

We next tested whether the putative sialic acid catabolic genes from *H. hathewayi* DSM13479 were sufficient to rescue growth on sialic acid in *E. coli* knockout strains from the Keio collection (58). As previously reported (29, 33, 50), *E. coli ΔnanA*, *ΔnanK*, and *ΔnanE* mutants failed to grow when cultured in minimal M9 medium with Neu5Ac as the sole carbon source, confirming the essential role of these genes in sialic acid metabolism. In contrast, all knockout strains, along with the wild-type strain, exhibited robust growth in minimal M9 medium supplemented with glucose, indicating that the observed growth defect is specific to the inability to utilize sialic acid (Fig. 5).

**Figure 5.**
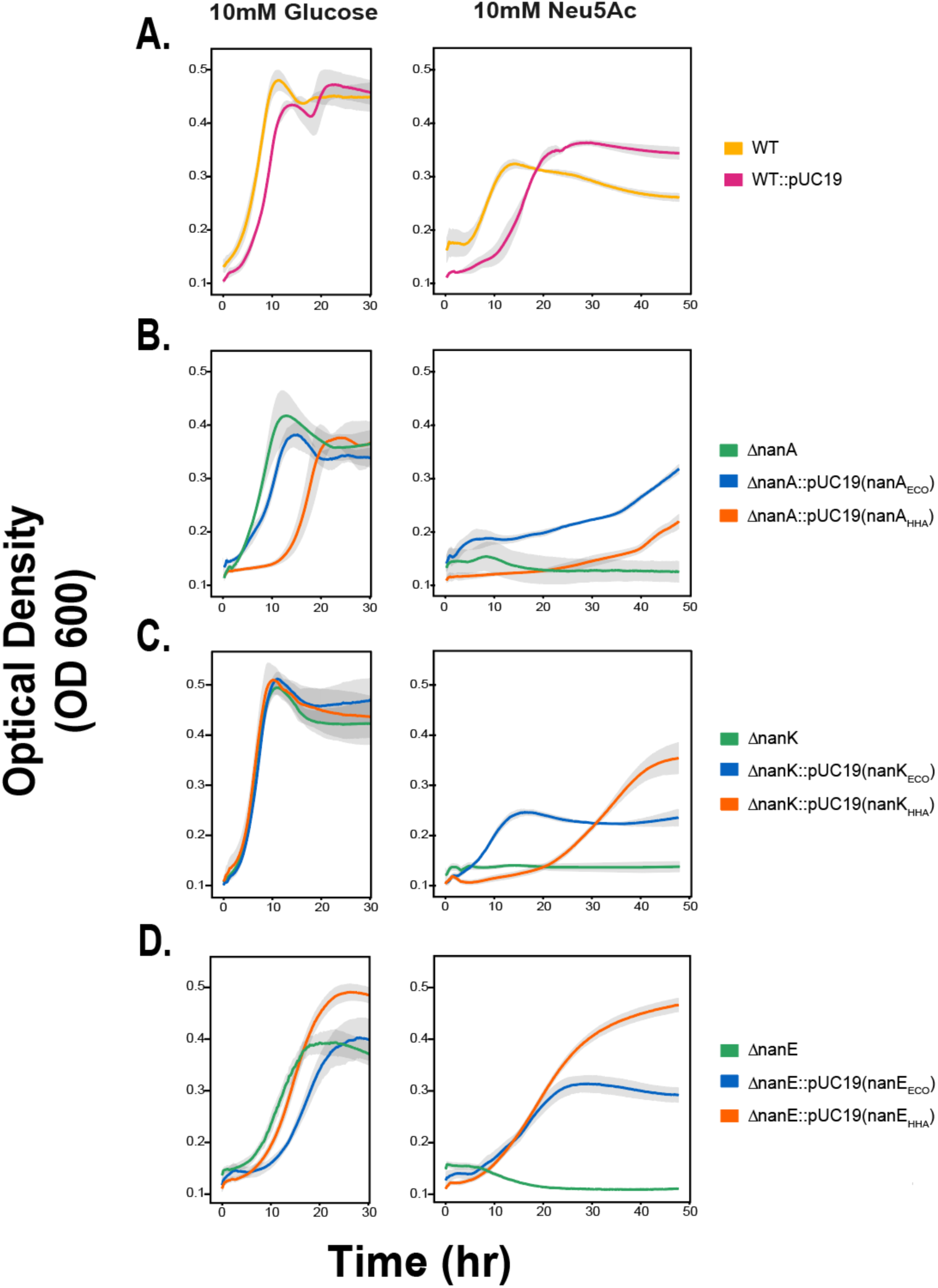
Growth of *E. coli* BW25113 wild type, *nanA/K/E* knockout (KO) strains, and complemented derivatives in glucose (left panels) and Neu5Ac (right panels). From top to bottom: **A.** wild type, **B.** *nanA* KO and complements, **C.** *nanK* KO and complements, and **D.** *nanE* KO and complements. Colors indicate genotype: wild type (yellow), wild type carrying empty pUC19 vector (pink), KO strains (green), KOs complemented with native *E. coli* genes (blue), and KOs complemented with the corresponding *H. hathewayi* genes (orange). [mean of 3 replicates +/- standard deviation].

To determine whether *H. hathewayi nan* orthologs are sufficient to restore sialic acid utilization in *E. coli* knockout strains, the corresponding *nanA, nanK,* and *nanE* genes were individually cloned into the IPTG-inducible expression vector pUC19 and introduced into the respective *E. coli* knockout strains. For comparison, the native *nan* genes from *E. coli* were also cloned and expressed under the same conditions. Expression of either the native *E. coli* or the *H. hathewayi nanA, nanK,* and *nanE* genes restored growth in each knockout strain (Fig. 5). This confirms that the *H. hathewayi* orthologs are functionally equivalent and capable of rescuing the metabolic defects of the *E. coli* knockouts. A control strain carrying the empty pUC19 vector (lacking any insert) was also included to evaluate the effect of plasmid carriage alone on bacterial growth. This strain exhibited a slightly extended lag phase compared to wild-type *E. coli* but ultimately reached a similar final optical density at the end of the exponential phase, indicating only a modest growth delay rather than a substantial fitness cost (Fig. 5A).

Although all complemented strains have the ability to utilize sialic acids, a notable difference was observed in the growth dynamics of the *nanA*-complemented strains (Fig. 5B). Interestingly, strains expressing *nanA* from *H. hathewayi* exhibited a consistently longer lag phase (∼22-24 hours) compared to those complemented with either the native *E. coli nanA* (∼12-14 hours) or the other *H. hathewayi* genes, *nanK* and *nanE* (∼14-16 hours and ∼10-12 hours, respectively) (Figs. 5C and 5D). This delayed onset of growth suggests that while the *H. hathewayi nanA* gene is functionally compatible, its enzymatic efficiency or regulation in the *E. coli* background might be suboptimal. In contrast, *nanK* and *nanE*, which encode a kinase and an epimerase respectively, did not show such delays (Figs. 5C and 5D), suggesting their enzymatic functions might be more promiscuous or less dependent on strict substrate recognition. Given that NanA is a lyase that must specifically recognize and cleave sialic acid, the observed lag could reflect a higher substrate specificity or reduced catalytic efficiency of the *H. hathewayi* NanA in the heterologous host. Further biochemical characterization could help determine whether these differences are due to enzyme kinetics, folding, or interactions with host-specific factors.

### H. hathewayi NanA shows functional conservation despite sequence divergence

Multiple sequence alignment and phylogenetic analysis of NanA amino acid sequences from the same 17 gut-associated *H. hathewayi* genomes previously selected from the BV-BRC database (Table S2) revealed that NanA is highly conserved within the strains showing pairwise similarities exceeding 98% (Fig. S3). Two additional BV-BRC genomes classified as *Hungatella spp.* with the same quality criteria were also included for genus-level comparison (Table S2). As shown in the phylogenetic tree (Fig. 6A), NanA protein sequences from nearly all *H. hathewayi* strains clustered together, forming a distinctive monophyletic group diverging from NanA sequences of reference organisms such as *E. coli*, *C. difficile*, and *Bacteroides* species, with the exception of strain HHA-3, whose truncated contig and orphan *nanA*-like annotation might reflect an assembly artifact rather than true phylogenetic divergence. Interestingly, the few *Hungatella*-associated NanA sequences that fell outside this main cluster correspond either to putative paralogs (e.g. second or third copies of *nanA* annotated within the same genome, Fig. S1), or to sequences found in fragmented genome assemblies (*i.e.* HHA-3), where annotation errors or misassemblies could not be ruled out. Notably, these outlier NanA sequences were often found in genomes that did not appear to encode a complete *nanAKE*-like pathway; despite containing additional *nanA* copies, these genomes lacked nearby annotations for *nanK*, or in some cases, both *nanK* and *nanE*. Additionally, one outlier originated from a *Hungatella sp*. strain not taxonomically classified as *H. hathewayi*, suggesting possible functional divergence within the genus. These observations support the idea that the primary *nanA* gene is evolutionarily conserved and likely functionally important across *H. hathewayi* strains, while additional or divergent copies may represent gene duplication events, assembly artifacts, or alternative metabolic configurations within the broader *Hungatella* lineage.

**Figure 6.**
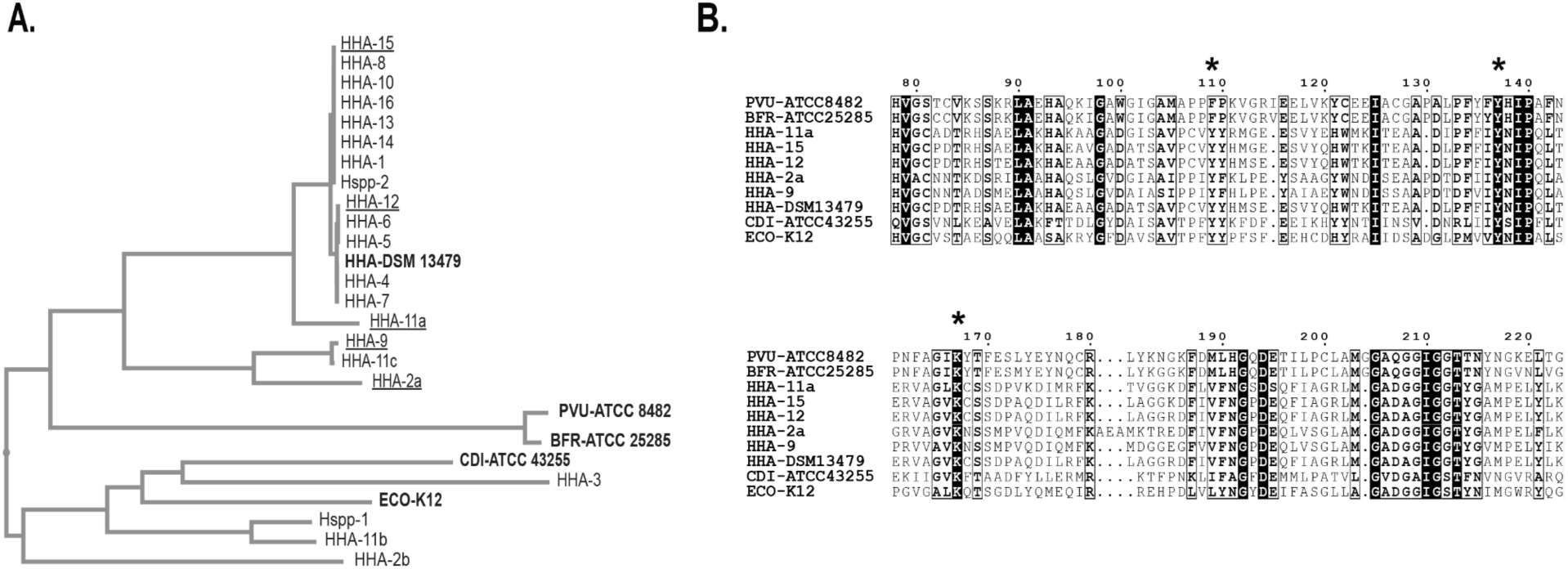
**A**. Phylogenetic tree of predicted NanA sequences from gut-associated *H. hathewayi* (HHA) and *Hungatella spp.* (Hspp) genomes, along with NanA sequences of reference organisms including *E. coli* K-12 (ECO)*, C. difficile* ATCC43255 (CDI), and *Bacteroides* species (*B. fragilis* ATCC25285 [BFR], *P. vulgatus* ATCC 8482 [PVU]). Strains underlined were selected as representatives of each cluster and serve as the sequences displayed in B. **B**. Multiple-sequence alignment (MSA) of representative NanA proteins illustrating with * strong conservation of catalytic residues Tyr111 (Y111), Tyr137 (Y137) and Lys165 (K165). Black-shaded boxes indicate positions where the amino acid is identical across all aligned sequences.

Despite the phylogenetic divergence of *H. hathewayi* NanA sequences from those of reference organisms such as *E. coli*, *C. difficile*, and *Bacteroides* species, the multiple sequence alignment reveals a high degree of conservation in key amino acid residues (Fig. 6B, Fig. S3). Residues associated with substrate binding and catalytic activity in *E. coli* NanA, previously identified through structural and biochemical studies (59–61), are fully conserved across all analyzed *H. hathewayi* NanA homologs. The multiple sequence alignment confirms strict conservation of Tyr137 (Y137) and Lys165 (K165), two residues shown to be essential for the formation of the Schiff-base intermediate and catalytic activity as demonstrated by crystallographic and mechanistic studies (59, 61). Lys165, in particular, has been identified as the nucleophilic residue responsible for Schiff-base formation with the sialic acid substrate, consistent with NanA classification as a class I aldolase. Notably, Tyr111 (Y111), which has been previously implicated in proton transfer and stabilization in *E. coli* NanA (60), is also conserved in NanA sequences from *H. hathewayi* strains. However, this residue is not conserved in more distantly related NanA homologs from reference species such as *B. fragilis* ATCC25285, or *P. vulgatus* ATCC8482, suggesting that while Y111 contributes to catalytic efficiency, it is not strictly essential for NanA function across all species.

## Discussion

Sialic acids such as Neu5Ac and Neu5Gc serve as critical nutrients in mucin-rich gut environments (17–19, 21). In this study, enrichment culturing from healthy human fecal microbiota under sialic acid supplementation revealed selection of *Hungatella* species across multiple donors, indicating a role for this genus as an active sialic acid degrader in the human gut. *H. hathewayi* genomes encode a conserved *nanAKE* gene cluster comprising the canonical enzymatic machinery for sialic acid catabolism (20, 30, 33, 39). We show that *H. hathewayi* can grow using either Neu5Ac or Neu5Gc as a sole carbon source suggesting the products of the NanA/K/E reactions can funnel into central carbon metabolism.

Sialic acid transport, regulation and catabolism genes in *H. hathewayi* are orthologous to genes encoded in an amalgam of phylogenetically distinct bacterial genomes. The *nanAKE* genes in *H. hathewayi* are orthologous to *E. coli* genes, yet the loci neighbors orthologs to genes involved in sialic acid metabolism in diverse bacterial species. Immediately downstream of *nanK* lies a ∼15 Kb region containing ABC-type transporter genes and an RpiR-family transcriptional regulator. RpiR-family regulators have been described in *Clostridium perfringens*, *Haemophilus influenzae*, *Vibrio vulnificus*, and *Staphylococcus aureus* (28, 52, 54, 55). ABC-type transport systems involved in sialic acid uptake have been described in *Haemophilus ducreyi*, *Streptococcus pneumoniae*, and *Bifidobacterium breve* (62–65). Thus uptake and regulation of sialic acids in *H. hathewayi* diverges from well-studied systems.

Growth of *H. hathewayi* equally on Neu5Ac and Neu5Gc means it can consume sialic acids derived from both host mucins and dietary meat. In contrast, *E. coli* grew faster on Neu5Ac than Neu5Gc, reflecting a preference for human-derived mucins. *H. hathewayi* reached higher maximum growth when sialic acids were used as a carbon source compared to glucose, while *E. coli* grew most on glucose. *H. hathewayi* appears to be a more specialized or efficient consumer, potentially adapted to niches where sialic acids are abundant.

Complementation of *E. coli ΔnanA*, *ΔnanK*, and *ΔnanE* mutants with *H. hathewayi* orthologs provides functional evidence that the predicted *nanAKE* cluster encodes active enzymes capable of restoring sialic acid metabolism. However, replacement of *E. coli nanA* with the *H. hathewayi* ortholog produced an extended lag phase, whereas *H. hathewayi nanK* and *nanE* enabled comparable growth to the *E. coli* orthologs. This observation is consistent with the precise substrate positioning required for Schiff-base formation and the sensitivity of lyases to subtle active-site variations (59–61, 66). Phylogenetic studies suggest that the lyase step represents a point of species-specific divergence within the Nan pathway (20). In contrast, the kinase and epimerase steps appear broadly tolerant across diverse bacteria, reflecting the conserved nature of their reactions and intermediates (20). Structural studies in NanK have shown highly conserved active sites as well as similar structural architecture among homologs from *Vibrio cholera, Pasteurella multocida* and *Haemophilus influenzae* (67). Likewise, structural and functional characterizations of NanE from organisms such as *Fusobacterium nucleatum*, *Vibrio cholerae*, and *Clostridium perfringens* underscore the conservation of 2-epimerization chemistry across bacterial lineages (35, 68) and the role of the ManNAc-6-P intermediate essential for central amino-sugar metabolism (20, 27). While the overall architecture of the Nan pathway is conserved across bacteria, our findings support that the lyase step (NanA) may diverge more than downstream reactions.

Phylogenetic analyses show that predicted NanA enzymes from *H. hathewayi* strains form a distinct clade separate from those of well-characterized homologs (*e.g.* NanA from *E. coli*, *C. difficile*, and *Bacteroides* spp.). Strong conservation of core catalytic residues, specifically Tyr111, Tyr137, and Lys165 underscores the evolutionary pressure to preserve enzymatic function despite sequence divergence. These specific residues have been experimentally validated in *E. coli* NanA to play essential roles in Schiff-base formation and proton transfer during sialic acid cleavage (59–61). Tyr111 is conserved in *H. hathewayi*, but not in more distantly related *Bacteroidota* species, suggesting it may be responsible for subtle differences in enzyme function rather than strictly required for activity.

*H. hathewayi* is a previously underappreciated participant in sialic acid consumption in the human gut. Probiotic modulation of sialic acid levels by species including *H. hathewayi* may prove useful to control inflammation and pathogen susceptibility in the gut. We used functional validation, comparative genomics, and phylogenetic analyses to show that *H. hathewayi* preserves core Nan pathway function while exhibiting lineage-specific divergence in enzyme behavior and loci structure. These initial steps to characterize the role of *H. hathewayi* in sialic acid metabolism will pave the way for a more thorough understanding of the role of *H. hathewayi* in host glycan utilization and gut microbial ecology.

## Acknowledgements

This work was supported by R35 GM147512 and an NIH supplement to Gabriel Miller, as well as an Amgen Scholarship awarded to Gabriel Miller. The funders had no role in study design, data collection and interpretation, or the decision to submit the work for publication. We thank the Titov Lab at University of California, Berkeley for access to LC–MS instrumentation and assistance with triple quadrupole mass spectrometry analyses. We thank Jessica Hoisington-Lopez and MariaLynn Crosby at Washington University in Saint Louis for sequencing support. We are grateful to the Deutschbauer Lab at University of California, Berkeley for providing *E. coli* strains and Keio knockout mutants used in this study.

## Materials and Methods

### Enrichment Culturing of Fecal Samples with Sialic Acids

#### Human Subjects and Ethical Approval

Fecal samples were collected from seven healthy adult donors under protocols approved by the Institutional Review Board (IRB) at the University of California (IRB protocol ID 2021-07-14511). All participants provided written informed consent prior to sample collection. Donors had not taken antibiotics within three months prior to donation. Fecal samples were collected using sterile collection kits (Medex Supply) and delivered to the lab within 24 hours of collection. Samples were stored on ice during transport and, upon arrival, were stored at –80 °C until further processing.

#### Enrichment Culture Setup

Fecal samples were pulverized using a pre-chilled mortar and pestle, with liquid nitrogen added throughout to maintain the sample in a frozen state during processing. Pulverized samples were then transferred into an anaerobic chamber (Coy Laboratory Products, 5% H₂, 20% CO₂, 75% N₂) and homogenized by vortexing with 2mm glass beads in phosphate-buffered saline containing 0.05% L-cysteine (PBS-Cys) at a 1:100 (w/v) dilution. The homogenates were filtered through a 100 µm cell strainer to obtain a clarified fecal suspension. Enrichment cultures were set up in deep-well plates by inoculating a defined base anaerobic medium (BAM; Table S1) with the clarified fecal suspension at a 1:100 dilution. BAM was supplemented with one of three sole carbon sources: 10 mM *N*-acetylneuraminic acid (Neu5Ac, Sigma), 10 mM *N*-glycolylneuraminic acid (Neu5Gc, Sigma), or 0.5% (w/v) porcine gastric mucin. Each condition was prepared in triplicate, along with controls lacking either inoculum or carbon source. Cultures were incubated anaerobically at 37°C for 48 hours. Following incubation, cultures were harvested for microbial community analysis via 16S rRNA gene sequencing.

#### Genomic DNA Extraction

Genomic DNA was extracted by bead-beating lysis followed by column purification (Monarch® gDNA Cleanup Kit, NEB). Culture pellets were transferred to 2 mL screw-cap tubes containing 500 µL 0.1 mm zirconia/silica beads, 500 µL Buffer A (200 mM Tris-HCl, pH 8.0; 200 mM NaCl; 20 mM EDTA), 210 µL 20% SDS, and 500 µL phenol/chloroform/isoamyl alcohol (25:24:1, pH 7.9). Samples were bead-beaten at high speed (2,400 rpm, BioSpec) for 4 min at room temperature, centrifuged (13,000 rpm, 3 min, 4 °C), and the aqueous phase was collected. The aqueous phase from each lysate was mixed with gDNA binding buffer (1:2, Monarch® gDNA Cleanup Kit, NEB), and binding and cleaning steps were performed according to the manufacturer’s instructions. Genomic DNA was eluted in nuclease-free water and stored at −20 °C until use. DNA purity and concentration were determined using a nanospectrophotometer (N50 NanoPhotometer, Implen).

#### 16S rRNA Amplicon Library Preparation

The V4 region of the 16S rRNA gene was amplified using the universal primers 515F (5′-GTGYCAGCMGCCGCGGTAA-3′) and 806R (5′-GGACTACNVGGGTWTCTAAT-3′), each with Nextera overhang adapters for downstream library preparation. DNA was amplified using two-step PCR (69). In the first PCR (PCR1), 10 ul reactions containing 2 µL of 5X KAPA HiFi buffer, 0.3 µL of 10 mM dNTPs, 0.5 µL DMSO, 0.2 µL of KAPA HiFi HotStart polymerase, 0.5 µL each of forward and reverse Nextera-tagged primers (10 µM), and 6 μL of 10ng/ul template DNA. Thermal cycling conditions were as follows: initial denaturation at 95 °C for 5 min; 25 cycles of 98 °C for 20 s, 55 °C for 15 s, and 72 °C for 1 min; with a final extension at 72 °C for 10 min. PCR1 amplicons were diluted 1:10 in nuclease-free water, and 5 µL of the diluted product was used as template in a second PCR (PCR2) to incorporate dual indexing primers. PCR2 (10 µL total volume) included 2 µL of 5X KAPA HiFi buffer, 0.3 µL of 10 mM dNTPs, 0.5 µL DMSO, 0.2 µL KAPA HiFi polymerase, 0.5 µL each of forward and reverse indexing primers (10 µM). Thermal cycling conditions were: 95 °C for 5 min; 10 cycles of 98 °C for 20 s, 55 °C for 15 s, and 72 °C for 1 min; and a final extension at 72 °C for 10 min. Indexed libraries were purified using magnetic bead-based size selection to remove primer dimers and other small fragments. A right-side size selection protocol using a 0.8:1 bead-to-sample ratio was performed according to the manufacturer’s instructions (SPRIselect beads, Beckman Coulter). To assess the size and integrity of the purified libraries (∼400 bp), the size-selected products were visualized on 1.0% agarose gels using electrophoresis. Final libraries were quantified using a Quanti-iT dsDNA High Sensitivity Kit (Thermo Fisher Scientific) according to the manufacturer’s instructions and pooled at equimolar concentrations for sequencing. Pooled libraries were assessed for quality and size distribution using 1.0% agarose gel electrophoresis and a BioAnalyzer (Agilent 2100) with the High Sensitivity DNA Kit (Agilent Technologies) following the manufacturer’s instructions.

#### Library Sequencing and Data Analysis

Sequencing was performed on the Illumina MiSeq platform using 2×250 bp paired-end chemistry with a MiSeq v2 500-cycle kit at the Edison Family Center for Genome Sciences & Systems Biology, Washington University in St. Louis. A 10% PhiX control library (Illumina) was spiked in to increase base diversity and serve as an internal sequencing control. Raw reads were processed and analyzed using QIIME 2 version 2022.8 (70) with the DADA2 pipeline for quality filtering, denoising, chimera removal, and inference of amplicon sequence variants (ASVs) (71). Taxonomic assignment was performed using a naïve Bayesian classifier trained on the SILVA reference database version 138.1 (72). Differential abundance analysis was conducted using ANCOM-BC to identify bacterial taxa significantly enriched under each substrate condition relative to baseline inoculum (73). Enriched taxa were interpreted as putative utilizers of the provided sialic acid or mucin substrate.

### Molecular Cloning of Hungatella Sialic Acid Genes into E. coli

#### Bacterial Strains and Growth Conditions

*Hungatella hathewayi* DSM 13479 was obtained from the DSMZ collection and cultured anaerobically in LYBHI medium (74), at 37°C in an anaerobic chamber (Coy Laboratory Products, 5% H₂, 20% CO₂, 75% N₂). The parental wild-type *E. coli* K-12 MG1655 strain and corresponding knockout strains lacking the sialic acid catabolic genes *nanA, nanK,* and *nanE* were obtained from the Keio collection (58). Aerobic cultures of *E. coli* were grown in Luria–Bertani (LB) broth or on LB agar plates supplemented with the appropriate antibiotics (100 µg/mL ampicillin or 50 µg/mL kanamycin). To assess sialic acid catabolism, 10 mM *N*-acetylneuraminic acid (Neu5Ac; Sigma) or 10 mM glucose (Sigma), as a control, was used as the sole carbon source in M9 minimal medium supplemented with 2 mM MgSO₄ and 0.1 mM CaCl₂.

#### Plasmid Design and Complementation of Knockout Strains

Plasmids were designed and synthesized by Twist Bioscience using the pUC19 backbone, with native sialic acid catabolic genes from *E. coli* and putative homologs from *H. hathewayi* cloned into the HindIII and XbaI restriction sites. Plasmids were transformed into the corresponding *E. coli* K-12 knockout strains (Δ*nanA*, Δ*nanK*, Δ*nanE*) via heat-shock transformation using in-house prepared chemically competent cells. Briefly, competent cells were prepared from single-colony cultures grown to mid-log phase (0.4–0.6 OD₆₀₀). Cell pellets were harvested and washed three times with ice-cold 100 mM CaCl₂. Following centrifugation and washing steps, pellets were resuspended in a minimal volume (∼100ul) of 100mM CaCl₂ with 15% (v/v) glycerol, resulting in an estimated ∼200X concentration relative to the original mid-log phase culture. After transformation, plasmids from overnight ampicillin-containing cultures were purified using the NEB Miniprep Kit (New England Biolabs) and sequence-verified by Nanopore sequencing (University of California Berkeley, DNA Sequencing Facility).

#### Growth Assays on Sialic Acid

Complemented knockouts, vector-only controls, and wild-type strains were grown overnight in LB broth supplemented with ampicillin (100 µg/mL) when required. Overnight cultures were washed twice in 1X PBS and resuspended to an OD₆₀₀ of 0.05 in M9 minimal medium containing either 10 mM Neu5Ac or 10 mM glucose as a growth control, along with ampicillin (100 µg/mL) and IPTG (1 mM). Cultures were set up in 96-well plates and incubated at 37 °C with shaking (200 rpm). Wells containing uninoculated media (with carbon source) and M9 minimal media without a carbon source were included as negative controls. OD₆₀₀ was recorded every 15 min for 48 h using a plate reader (Accuris^TM^ SmartReader^TM^ 96 Microplate Absorbance Reader). OD data were processed and plotted using R version 2025.09.1. Growth curves were analyzed to assess functional complementation.

### LC-MS Quantification of N-acetylneuraminic acid

#### Metabolite Extraction

Culture supernatants from *E. coli* and *H. hathewayi* were collected at two time points: immediately after inoculation and upon reaching stationary phase (OD ∼0.6). *E. coli* was grown aerobically in LB medium, while *H. hathewayi* was grown anaerobically in LYBHI medium. Cultures were grown in triplicate. At each time point, culture aliquots were centrifuged to pellet the cells, and the resulting supernatants were immediately frozen and stored at −80 °C for subsequent metabolite extraction. Metabolites were extracted from 2 μL of culture supernatant by adding 998 μL of pre-chilled extraction buffer, consisting of 400 μL methanol, 400 μL acetonitrile, 192.15 μL HPLC-grade water, 3.85 μL 98% formic acid, and 1 μL each of 1 mM stable isotope-labeled internal standards (glucose-6-phosphate and fructose-6-phosphate). Extracts were vortexed and centrifuged at 17,000 g for 10 min at 4 °C. Then, 900 μL of the supernatant extract was neutralized with 100 μL of pre-chilled 1M ammonium bicarbonate. Neutralized extracts were aliquoted and stored at −80 °C until LC-MS analysis.

#### LC-MS Setup and Analysis

LC-MS analysis was performed using an iHILIC-(P) Classic column (PEEK, 200 Å, 5 μm, 2.1×150 mm; Analytics-Shop USA LP) on an Agilent 6430 Triple Quadrupole mass spectrometer operated in negative ion mode. The mobile phase consisted of 100% LC-MS grade acetonitrile as organic phase and 20 mM ammonium carbonate with 0.1% ammonium hydroxide as aqueous phase. Flow rate was set to 0.150 mL/min. Each sample (20 μL injection) was run for 1h. A standard curve prepared with known concentrations of Neu5Ac was used to quantify Neu5Ac in the samples. Data were acquired and analyzed using Agilent MassHunter Quantitative Analysis software version 3.6.

