## Supplemental materials for "A bidirectional *nanAKE* locus enables sialic acid catabolism in gut microbiome member *Hungatella hathewayi*"

### Supplemental Material

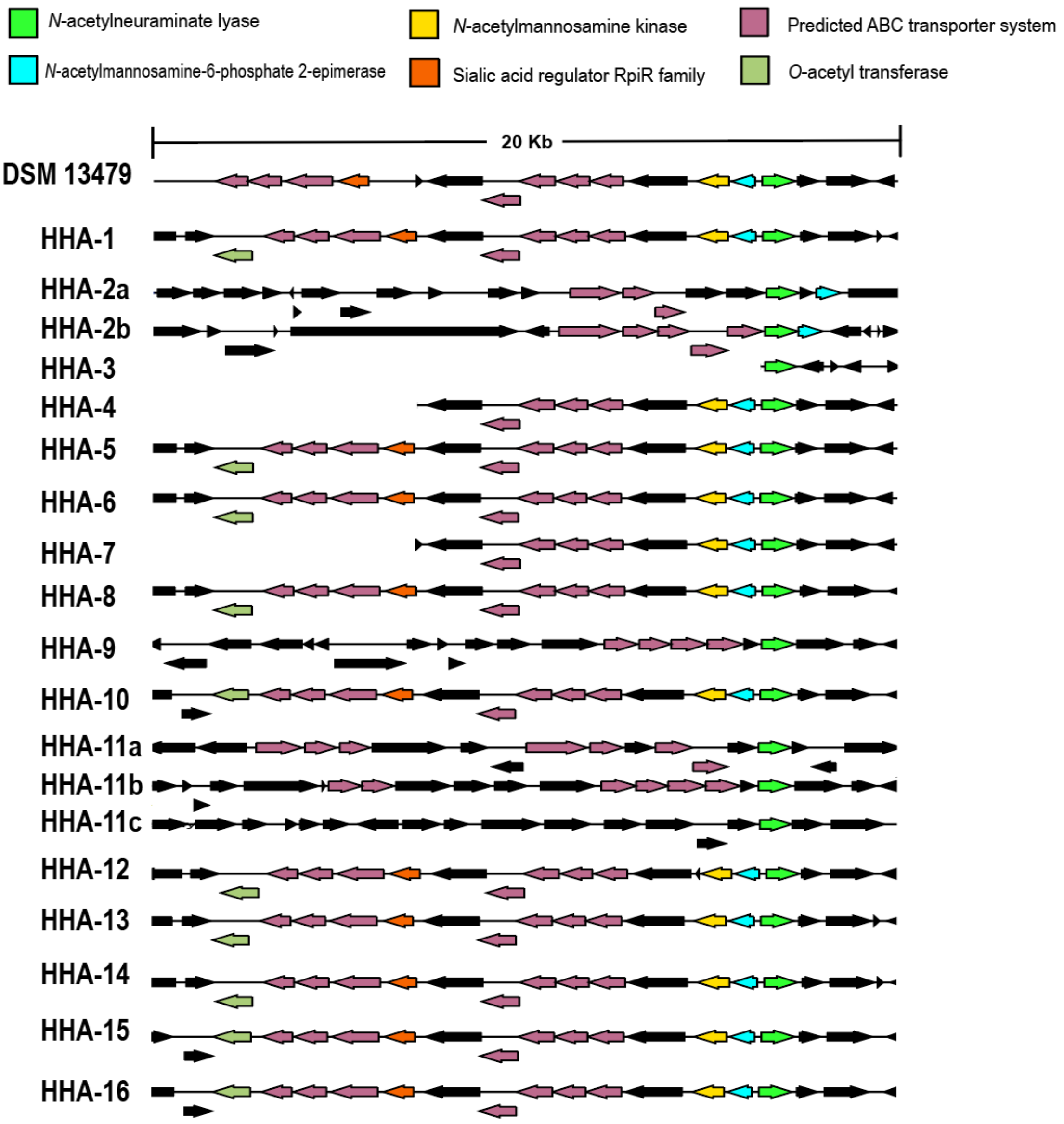


**Figure S1.** Putative sialic acid-catabolic genes annotated in publicly available genomes of gut-associated *Hungatella hathewayi* strains including the type strain *H. hathewayi* DSM13479. Genes are represented as arrows indicating orientation and scaled to their relative lengths.

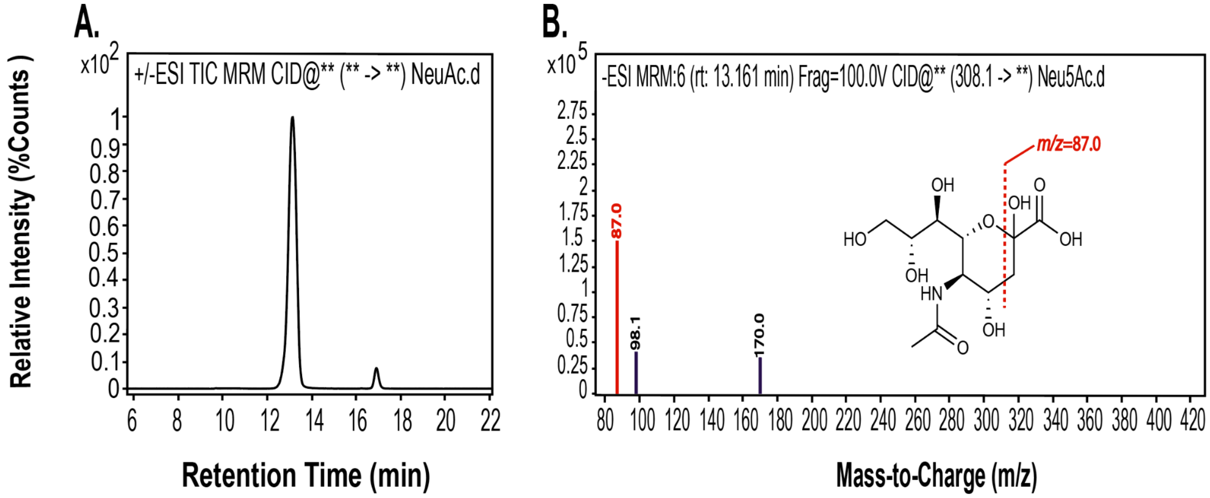


**Figure S2.** LC-MS/MS analysis of *N*-acetylneuraminic acid (Neu5Ac) using Multiple Reaction Monitoring (MRM). **A.** Extracted chromatogram showing a major peak at ~13.2 min, corresponding to the retention time of Neu5Ac under our chromatographic conditions. **B.** Product ion (MS/MS) spectrum acquired at ~13.2 min for the MRM transition m/z 308.1 (precursor) to fragment ions, indicating the fragmentation pattern of Neu5Ac in negative electrospray ionization (ESI) mode. *m/z* 87.0 was selected for quantification.

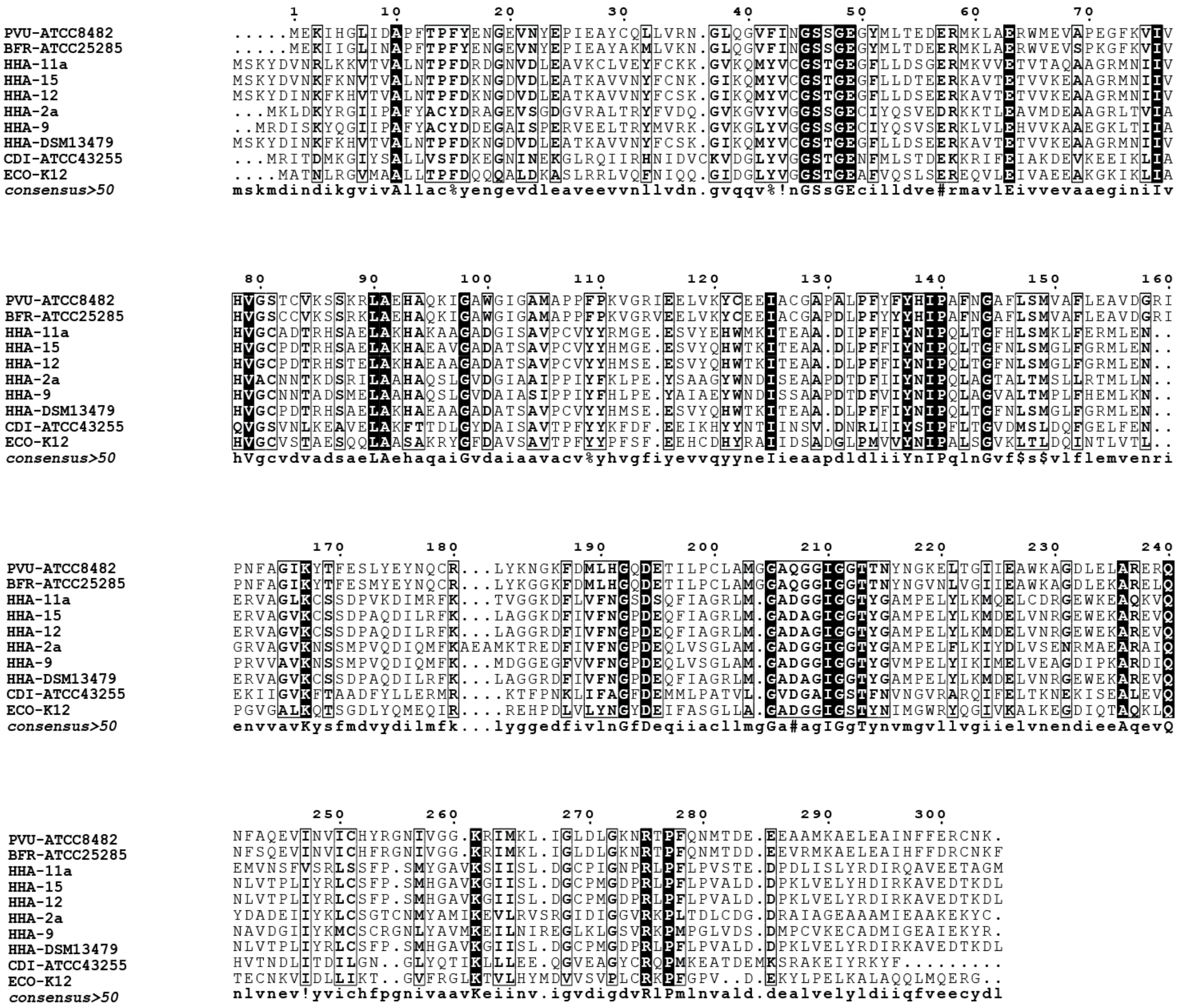

**Figure S3.** Full-length multiple sequence alignment (MSA) of NanA amino acid sequences from representative *H. hathewayi* (HHA) and *Hungatella* spp. (Hspp) genomes, along with reference sequences from *E. coli* K-12 (ECO), *C. difficile* ATCC43255 (CDI), *B. fragilis* ATCC25285 (BFR), and *P. vulgatus* ATCC8482 (PVU). This alignment corresponds to the complete NanA protein sequences underlying the partial alignment shown in Fig 5B, where conserved catalytic residues were highlighted. Black-shaded boxes indicate positions where the amino acid is identical across all aligned sequences.

**Table S1.** Composition of Base Anaerobic Media (BAM). List of each ingredient with the corresponding final concentration in the prepared medium

| Component | Concentration |
|---|---|
| L-Cysteine·HCl | 0.5 g/L |
| $KH_2PO_4$ | 4.1 g/L |
| $K_2HPO_4$ | 12.2 g/L |
| $MgSO_4 \cdot 7H_2O$ | 0.02 g/L |
| $NaHCO_3$ | 0.4 g/L |
| NaCl | 0.08 g/L |
| $CaCl_2$ | 8 mg/ml |
| $FeSO_4 \cdot 7H2O$ | 0.4 mg/ml |
| Menadione | 1 mg/L |
| Hematin | 1.2 mg/L |
| L-Histadine | 31 mg/L |
| Tween-80 | 0.5 ml/L |
| ATCC Mineral Mix | 10 ml/L |
| ATCC Vitamin Mix | 10 ml/L |

**Table S2.** *nanA* annotations used in this study. Columns list this study-assigned ID, source genome, genome ID, NCBI accession number, BV-BRC ID, and RefSeq locus tag corresponding to each *nanA* annotation.

| This study ID | Source genome | Genome ID | NCBI Accession# | BRC ID | RefSeq Locus Tag |
|---|---|---|---|---|---|
| ECO-K12 | Escherichia coli str. K-12 substr. MG1655 | 511145.12 | NC_000913 | fig\|511145.12.peg.3321 | b3225 |
| BFR-ATCC 25285 | Bacteroides fragilis NCTC 9343 strain ATCC 25285 | 272559.17 | NC_003228 | fig\|272559.17.peg.1672 | BF1712 |
| PVU-ATCC 8482 | Bacteroides vulgatus ATCC 8482 | 435590.9 | NC_009614 | fig\|435590.9.peg.4228 | BVU_4117 |
| CDI-ATCC 43255 | Clostridium difficile ATCC 43255 | 499175.4 | NZ_CM000604 | fig\|499175.4.peg.1867 | CdifA_020200011752 |
| HHA-DSM 13479 | Clostridium hathewayi DSM 13479 | 566550.8 | NZ_GG667621 | fig\|566550.8.peg.1224 | |
| HHA-1 | Hungatella hathewayi strain AF19-13AC | 154046.46 | QTJW01000012 | fig\|154046.46.peg.1051 | DWX31_18190 |
| HHA-2a | Hungatella hathewayi strain AF31-1 | 154046.43 | QVHZ01000002 | fig\|154046.43.peg.2466 | DWZ21_06175 |
| HHA-2b | Hungatella hathewayi strain AF31-1 | 154046.43 | QVHZ01000003 | fig\|154046.43.peg.3987 | DWZ21_12535 |
| HHA-3 | Hungatella hathewayi strain AM21-16 | 154046.45 | QUSI01000018 | fig\|154046.45.peg.1142 | DW241_11855 |
| HHA-4 | Hungatella hathewayi strain AM39-16AC | 154046.53 | QSGX01000065 | fig\|154046.53.peg.6051 | DW876_30770 |
| HHA-5 | Hungatella hathewayi strain CE91-St54 | 154046.356 | BQNK01000001 | fig\|154046.356.peg.6314 | CE91St54_59450 |

| | | | | | |
|---|---|---|---|---|---|
| HHA-6 | Hungatella hathewayi strain CE91-St55 | 154046.355 | BQNJ01000001 | fig\|154046.355.peg.3550 | CE91St55_33410 |
| HHA-7 | Hungatella hathewayi strain L1_007_061G1_dasL1_007_061G1_metabat.metabat.9 | 154046.305 | JAGZVY010000096 | fig\|154046.305.peg.3864 | KH325_23935 |
| HHA-8 | Hungatella hathewayi strain L3_129_062G1_dasL3_129_062G1_maxbin2.maxbin.021sta_sub | 154046.304 | JAGZHP010000004 | fig\|154046.304.peg.1205 | KHZ57_05450 |
| HHA-9 | Hungatella hathewayi strain L3_131_368G1_dasL3_131_368G1_metabat.metabat.46 | 154046.301 | JAGZLN010000031 | fig\|154046.301.peg.2421 | KH024_20160 |
| HHA-10 | Hungatella hathewayi strain L3_133_000G1_dasL3_133_000G1_metabat.metabat.26 | 154046.302 | JAGZKD010000004 | fig\|154046.302.peg.1592 | KH010_04445 |
| HHA-11a | Hungatella hathewayi strain L3_133_000M1_dasL3_133_000M1_metabat.metabat.43 | 154046.303 | JAGZKF010000004 | fig\|154046.303.peg.2390 | KHZ58_02595 |
| HHA-11b | Hungatella hathewayi strain L3_133_000M1_dasL3_133_000M1_metabat.metabat.43 | 154046.303 | JAGZKF010000020 | fig\|154046.303.peg.1615 | KHZ58_07860 |
| HHA-11c | Hungatella hathewayi strain L3_133_000M1_dasL3_133_000M1_metabat.metabat.43 | 154046.303 | JAGZKF010000032 | fig\|154046.303.peg.2575 | KHZ58_10495 |
| HHA-12 | Hungatella hathewayi strain MCC342 | 154046.308 | WQPR01000039 | fig\|154046.308.peg.4902 | GPL27_15645 |
| HHA-13 | Hungatella hathewayi strain OM02-1 | 154046.49 | QSVU01000013 | fig\|154046.49.peg.1204 | DXB08_15945 |
| HHA-14 | Hungatella hathewayi strain TF05-11AC | 154046.48 | QSSQ01000029 | fig\|154046.48.peg.3436 | DXC39_22475 |

| | | | | | |
|---|---|---|---|---|---|
| HHA-15 | Hungatella hathewayi strain TF09-11AC | 154046.47 | QSRE01000002 | fig\|154046.47.peg.3001 | DXC88_05325 |
| HHA-16 | Hungatella hathewayi strain TM09-12 | 154046.51 | QSON01000006 | fig\|154046.51.peg.5995 | DXD79_13855 |
| Hspp-1 | Hungatella sp. L12 | 2763050.3 | JACOPB010000002 | fig\|2763050.3.peg.1672 | H8S75_07845 |
| Hspp-2 | Hungatella sp. L36 | 2763049.3 | JACOPC010000003 | fig\|2763049.3.peg.2125 | H8S80_09755 |

**Table S3.** Log-phase growth rates and lag phases of *H. hathewayi* DSM13479 and *E. coli* BW25113 on different carbon sources. Values represent mean ± standard deviation from three biological replicates. Growth rates were calculated from the slope of the linear region of log-transformed OD600 values during exponential phase. Lag phase was estimated as the x-intercept of the extrapolated log-phase regression line.

| Organism | Carbon source | Log-phase growth rate ($hr^{-1}$) | Lag phase (hr) |
|---|---|---|---|
| *H. hathewayi* | Glucose | 0.0263 ± 0.004 | 12.06 ± 0.38 |
| | Neu5Ac | 0.0427 ± 0.0031 | 13.17 ± 0.14 |
| | Neu5Gc | 0.0435 ± 0.0041 | 13.50 ± 0.29 |
| *E. coli* | Glucose | 0.277 ± 0.013 | 1.94 ± 0.13 |
| | Neu5Ac | 0.091 ± 0.001 | 8.02 ± 0.4 |
| | Neu5Gc | 0.0368 ± 0.0017 | 16.02 ± 0.2 |
